# Potential cross-protection against SARS-CoV-2 from previous exposure to bovine coronavirus

**DOI:** 10.1101/2021.12.13.472476

**Authors:** Lana Bazan Peters Querne, Fernanda Zettel Bastos, Breno Castello Branco Beirão

**Affiliations:** Programa de Pós Graduação em Engenharia de Bioprocessos e Biotecnologia, Universidade Federal do Paraná. Av. Cel. Francisco H. dos Santos, 100, Centro Politécnico, Curitiba, PR, Brazil, CEP 81531-980; Departamento de Patologia Básica, Universidade Federal do Paraná. Av. Cel. Francisco H. dos Santos, 100, Setor de Ciências Biológicas, Curitiba, PR, Brazil, CEP 81531-980

**Keywords:** bovine coronavirus (BCoV), epitope, in silico, SARS-CoV-2

## Abstract

Humans have long shared infectious agents with cattle, and the common cold OC-43 CoV is a not-so-distant example of cross-species viral spillover. Human exposure to BCoV is certainly common, as the virus is endemic in cattle-raising regions. This article shows an *in silico* investigation of shared viral epitopes between BCoV and SARS-CoV-2. HLA recognition and lymphocyte reactivity were assessed using freely-available resources. Several epitopes were shared between BCoV and SARS-CoV-2, both for B and T lymphocytes. These data demonstrate that possible cross-protection is being induced by human exposure to cattle.

## 1. Introduction

In December 2019, a new coronavirus named severe acute respiratory syndrome coronavirus 2 (SARS-CoV-2) was discovered in Wuhan, China’s Hubei province [1]. The SARS-CoV-2 is responsible for coronavirus disease 2019 (COVID-19), characterized by an unusual pneumonia outbreak that led the World Health Organization to declare the new coronavirus pandemic on March 11, 2020. Until October 11, 2021, COVID-19 was responsible for more than 237 million cases and more than 4.8 million deaths around the world [2].

The symptoms of infected people resemble those of viral pneumonia, such as cough, fever and discomfort when breathing [3]. In elderly patients and patients with comorbidities (e.g. diabetes, obesity and asthma) the development of severe cases with dyspnea and bilateral pulmonary infiltration is more common, increasing the number of hospitalizations and deaths in this population [4].

Coronaviruses are single-strand RNA viruses belonging to the *Coronaviridae* family, capable of infecting several animals causing respiratory, gastrointestinal and neurological diseases. The four genera that compose this family are: *Alphacoronavirus*, *Betacoronavirus*, *Gammacoronavirus* and *Deltacoronavirus* [5]. Among the beta-coronaviruses are the aforementioned SARS-CoV-2 and the bovine coronavirus (BCoV), being the latter responsible for livestock losses causing diarrhea in newborn calves and respiratory infections in calves and confined cattle [6, 7]. Both viruses encode four structural proteins in the genome: envelope protein (E), membrane protein (M), nucleocapsid protein (N) and spike protein (S); in addition to non-structural proteins (NSP) and open reading frame polyproteins (ORF) [8, 9].

Cross-reactivity between coronavirus strains may be able to induce an adaptive immunity that would help reduce the severity and spread of disease. This pre-existing immunity from contact with other coronaviruses, such as BCoV, possibly can provide protection against the SARS-CoV-2 [10]. Nevertheless, previous works predicting cross-immunity have focused on coronaviruses from different genera [11].

As adaptive immunity is generated by the recognition of epitopes by T and B cells [12], the present study aimed to search for peptides originated from BCoV proteins M, N, S and ORF that presented T and B cell responses in humans and high identity with SARS-CoV-2.

## 2. Methods

### 2.1 Setting-up the peptides for analysis

The proteome sequences of bovine coronavirus were obtained from the NCBI database and focused on four proteins (Table 1): spike protein, membrane protein, nucleocapsid protein and replicase polyprotein (Orf1ab). The entire protein sequences were organized in 15-mer peptides that overlapped by 10 amino acids.

**Table 1.** Proteome sequences of bovine coronavirus and human SARS-CoV-2 obtained from NCBI. Accession numbers are indicated in the table.

|  | **Bovine coronavirus** | **SARS-CoV-2** |
| --- | --- | --- |
| Spike protein | NP_150077.1 | YP_009724390.1 |
| Membrane protein | NP_150082.1 | YP_009724393.1 |
| Nucleocapsid protein | NP_150084.1 | QQD86936.1 |
| Orf1ab | NP_150073.3 | BCT04066.1 |

Orf1ab = replicase polyprotein.

### 2.2 Prediction of T Cell Reactivity

T cell reactivity of bovine coronavirus peptides was assessed by predicting their binding to human leukocyte antigen class II (HLA II) molecules using IEDB MHC II binding predictions tool (http://tools.iedb.org/mhcii/). Peptide binding was predicted to all HLA class II molecules. A 20% percentile rank cutoff was chosen as a universal prediction threshold [13].

### 2.3 Prediction of B Cell Reactivity

B cell reactivity of bovine coronavirus peptides was assessed using IEDB Bepipred Linear Epitope Prediction 2.0 (http://tools.iedb.org/bcell/). The residues with scores above the threshold (0.5) and with 5 amino acids or more were predicted to be part of an epitope.

### 2.4 Identity of bovine peptides with human SARS-CoV-2 proteins

All bovine peptides that were above the threshold for T cells and B cells were analyzed for identity to the corresponding proteins of human SARS-CoV-2 (Table 1) using the Multiple Sequence Alignment (Clustal Omega, https://www.ebi.ac.uk/Tools/msa/clustalo/). Sequences with an identity greater than or equal to 80% were selected as peptide matches [14].

### 2.5 Epidemiology of COVID-19 and association with risk factors

COVID-19 epidemiology was assessed from publicly available data [15]. The slope of increase of cases/100,000 people for each city in the Brazilian State of Mato Grosso do Sul (MS) was used (between January, 2020 and September, 2021) [16]. The slope of COVID-19 cases was compared to the number of cattle/100,000 people for each municipality in the state [17].

As a control, the distance from each municipality to the major city in the subregion of the state was compared to the slope of COVID-19 cases [18]. General efficiency of public spending (not directly correlated with COVID-19) was also used as a control in a correlation analysis with COVID-19 prevalence. Data from the literature on public investment were used. Spending rigour was scored from 1-4, with four being the best-quality public use of resources [19]. The correlation of the data with COVID-19 prevalence was assessed with run’s test in a linear correlation. GraphPad Prism 8 (GraphPad Software, Inc., USA) was used for graphing and for statistical analysis. All the data used for this analysis is available as supplementary material (Supplementary Table 1).

## 3. Results

### 3.1 Setting up the Peptides for Analysis

A total of 136, 23, 45 and 709 15-mer peptides that overlapped by 10 amino acids were obtained for proteins S, M, N and ORF1ab respectively.

### 3.2 Prediction of T Cell Reactivity

From the results obtained by the IEDB MHC II binding prediction tool, 106 peptides from protein S, 20 peptides from protein M, 24 peptides from protein N and 566 peptides from ORF1ab protein had a percentile rank equal to or less than 20%.

### 3.3 Prediction of B Cell Reactivity

From the results obtained by the IEDB Bepipred Linear Epitope Prediction 2.0, 70 peptides from protein S, 9 peptides from protein M, 38 peptides from protein N and 386 peptides from ORF1ab protein had scores above the threshold.

### 3.4 Identity of bovine peptides with human SARS-CoV-2 proteins

Among the peptides that showed good results for T or B cells, only 2 peptides from protein S, 1 peptide from protein M, and 2 peptides from protein N showed at least 80% similarity with SARS-CoV-2 (Table 2). For these three proteins, no sequence was found to be within the cutoff values for both T cells and B cells.

**Table 2.**
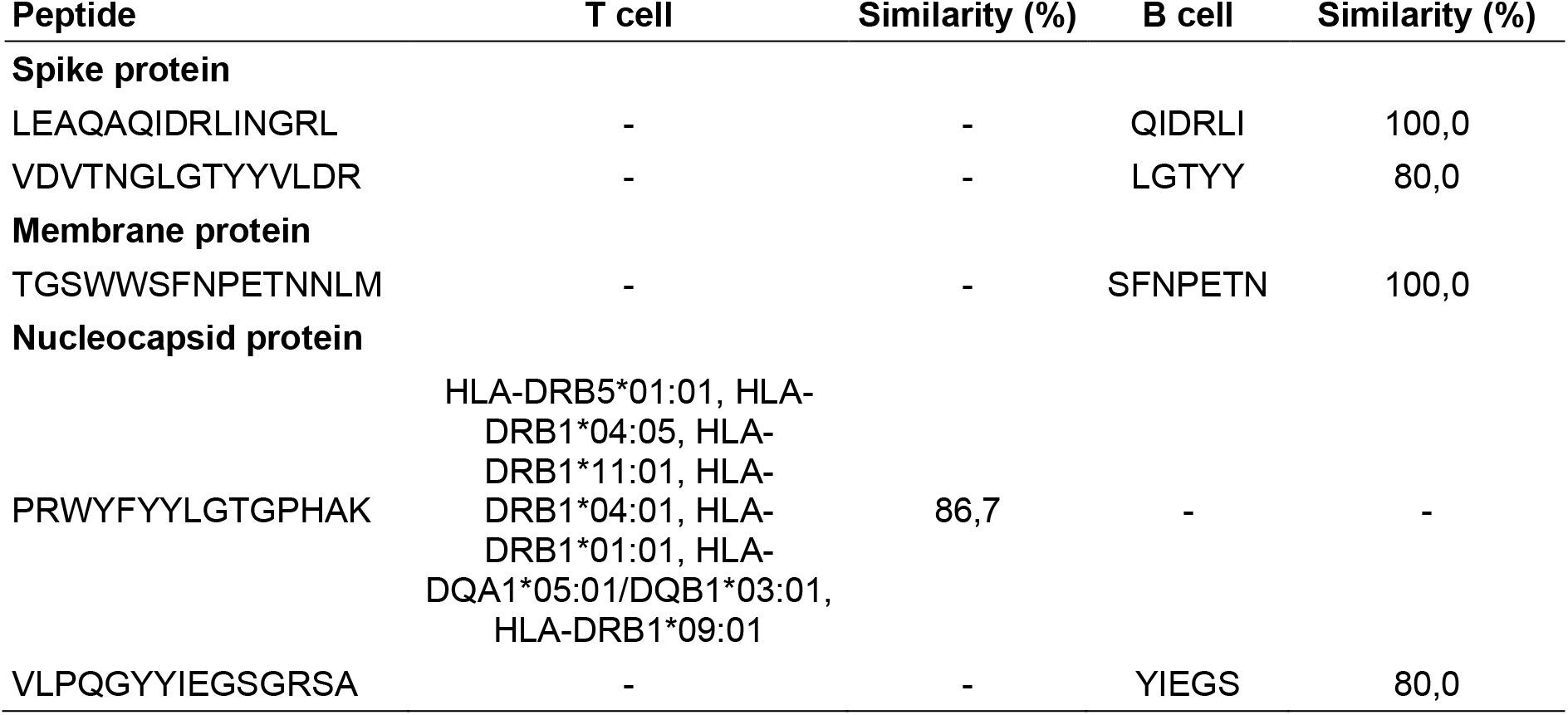
Peptides from Spike, membrane and nucleocapsid proteins that showed at least 80% similarity with SARS-CoV-2.

Regarding the ORF1ab protein,107 peptides showed good results for T or B cells (Table 3). In this case, 28 peptides were found to be within the cutoff values for both T cells and B cells.

**Table 3.**
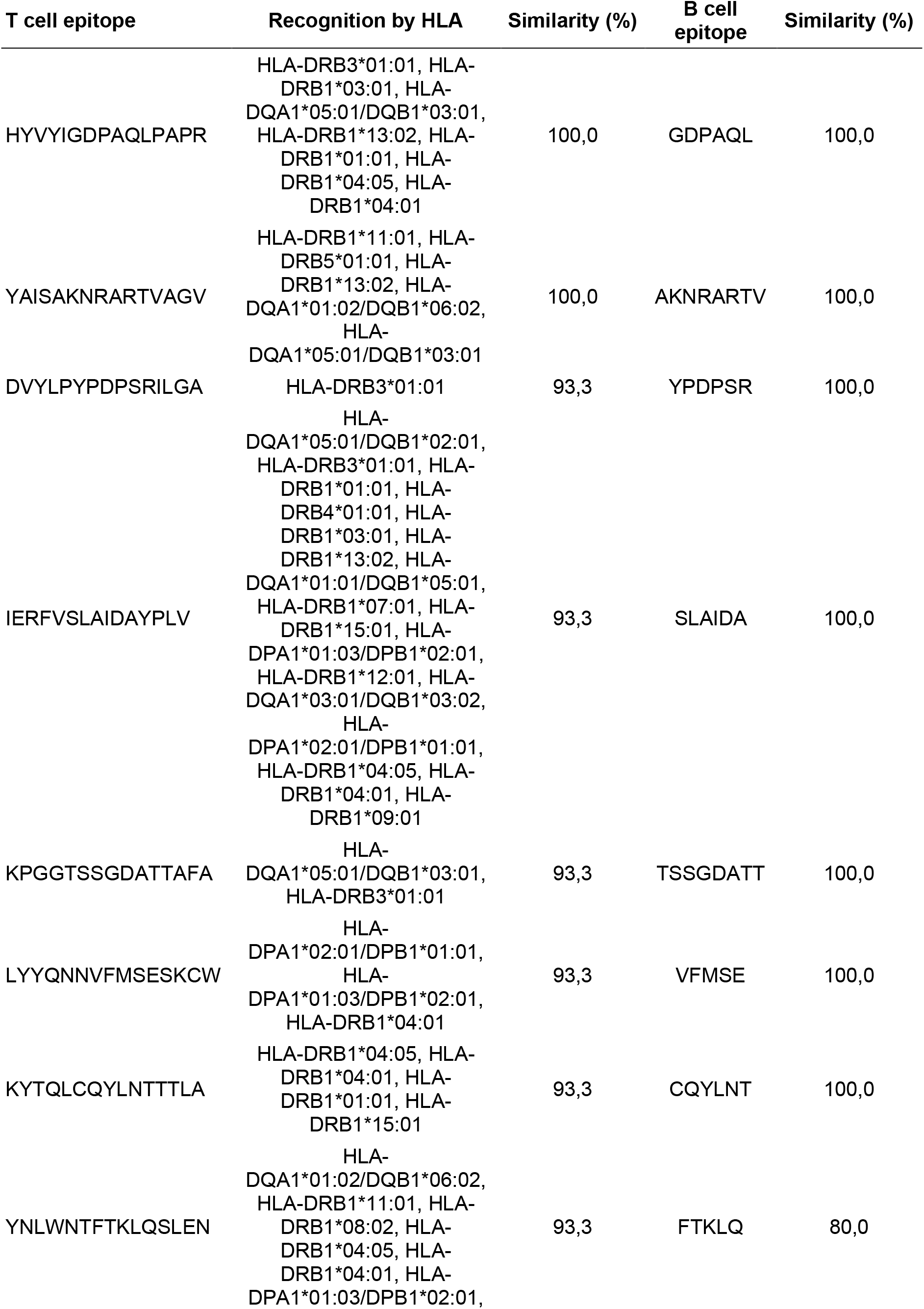
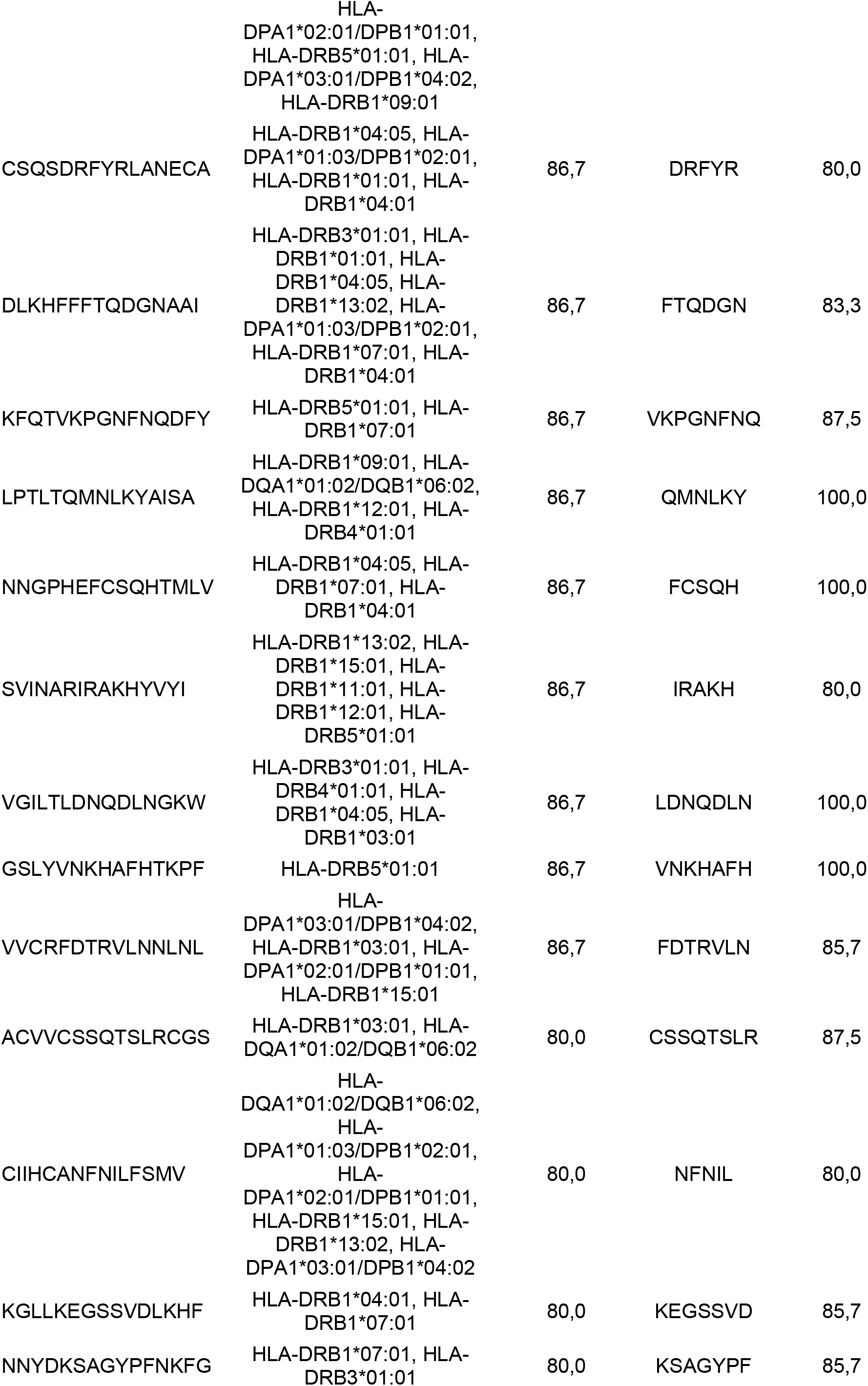
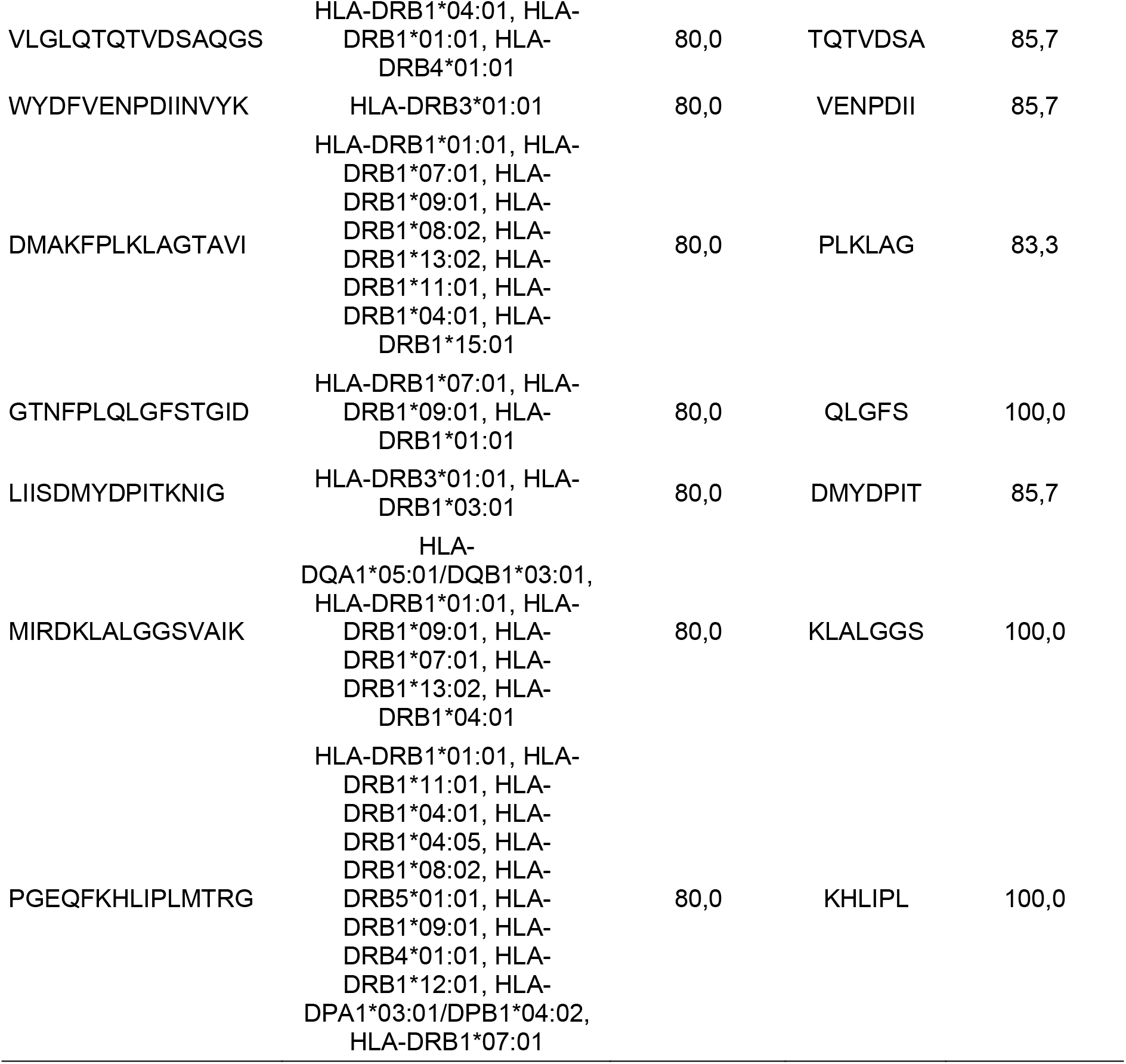
Peptides from replicase polyprotein (ORF1ab) that showed at least 80% similarity with SARS-CoV-2.

### 3.5 Epidemiology of COVID-19 and Association with Risk Factors

We analysed the correlation of COVID-19 prevalence to the density of cattle in the Brazilian state of MS as an example of a possible epidemiological association between human exposure to the Bovine Coronavirus (BCoV) and altered pandemic spread.

Cattle density (cattle/100,000 people) negatively correlated with the slope of COVID-19 case increase in MS. In opposition, confounding factors in this epidemiological analysis showed no association with the slope of COVID-19 cases in the state (assessed factors were distance of each municipality to the main regional hub city and quality of public spending) (Fig. 1).

**Figure 1.**
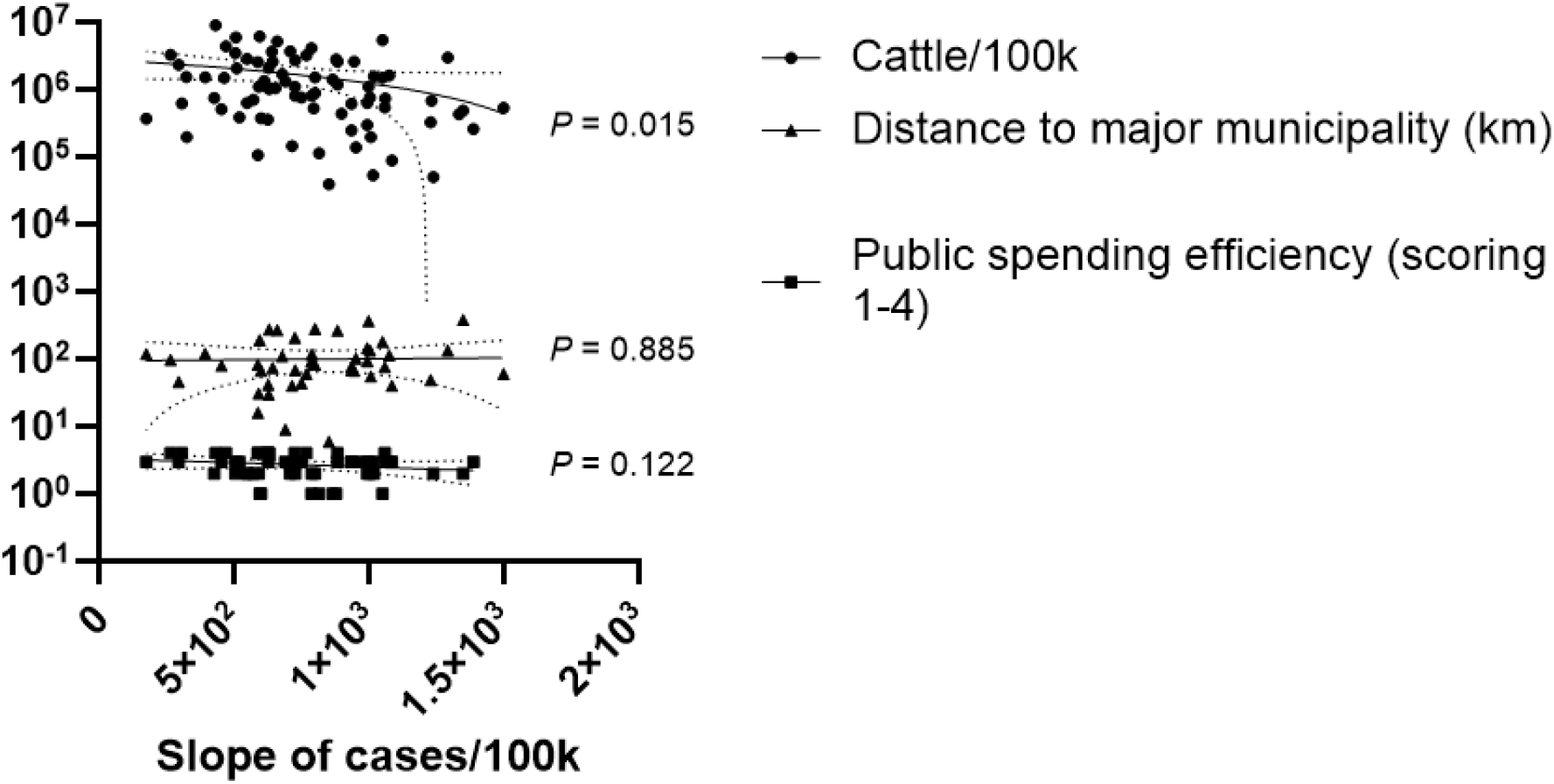
Linear regression between cattle density and the slope of cumulative COVID-19 case increase in the Brazilian State of Mato Grosso do Sul. Data between Jan/20 and Sep/21 were used. Cattle density was calculated as the number of cattle/100,000 people in the municipality. The distance of the municipality to the major hub city was used to control for lower people connectivity of cattle-raising areas. Public spending efficiency was used to control for possible slower responses to the COVID-19 pandemic from cattle-raising municipalities. Analysis by run’s test in a linear regression. *P-*values are shown for each regression. The dotted lines around the linear regression trend indicate the 99% CI.

## 4. Discussion

The premise of this article is the hypothesis that if BCoV influenced human immune responses to COVID-19, the epidemiology of the pandemic would have been altered by human exposure to cattle, since BCoV naturally occurs in bovine herds [20], in addition to other factors that can influence COVID-19 geographic spread, such as income rates and social vulnerability levels [21].

The Brazilian state of MS was chosen as an hypothesis-raising case study, as it is a large beef productor with no megacities, which can “distort” the local epidemiological status due to their large influence on the statistics and their worldwide connections [22, 23]. Within-state infrastructure, scholarity, income and animal production conditions are more homogeneous than in inter-state comparisons, and a whole-country analysis would be biased by these and therefore was not performed [24, 25]. Further, general efficiency of public spending was an important factor in the spread and control of the pandemic in Brazil [26], being another discrepant factor in the analysis of the whole-country.

Municipalities with more cattle are expected to be further away from regional hubs, since large land areas are needed for farming. Therefore, any association between COVID-19 cases with cattle density could only indicate lower connectivity of the municipality, which is a major cause of spatial proliferation of the disease [27].

Bovine coronaviruses (BCoV) are members of the *Betacoronavirus* genus together with SARS-CoV-2, denoting their similarities. Further, within the *Betacoronavirus*, BCoV is among the most similar to SARS-CoV-2 [28, 29]. Indeed, cattle can be experimentally infected with SARS-CoV-2 [30] and bovine coronaviruses have spilled over to humans before - current strains of BCoV can be cultured in human rectal adenocarcinoma cells, demonstrating that cross-species infection is still a risk, if not a common event already [31 – 33]. Other works have already discussed the immunological impacts that coronaviruses of domestic animals could have on humans. In Brazil, the use of the *Deltacoronavirus* Avian Infectious Bronchitis is being clinically tested for COVID-19 vaccination, for instance [11, 29]. Nevertheless, such considerations have not yet been directed to BCoV.

It was not the goal of this study to confirm the association of BCoV with COVID-19 using epidemiological data. Our analysis is exceedingly restricted for this purpose. We used epidemiological data of COVID-19 for hypothesis-raising, and they showed a possible association of human exposure to cattle with the development of the pandemic.

This work evaluated in silico if BCoV epitopes could be recognized by human B and T lymphocytes. Here, we report several epitopes which are likely to be important in the response for COVID-19 and which are shared with BCoV. This analysis is valuable in understanding the impact that exposure to the bovine coronavirus may have had on COVID-19. It is possible that COVID-19 epidemiology was shaped by human exposure to BCoV, much as smallpox was naturally curtailed by the exposure to cowpox [34], and this paper offers evidence for viral cross-protection from the analysis of common epitopes.

## 5. Conclusion

SARS-CoV-2 and BCoV share several common epitopes, which may confer cross-immunity. The relevance of this connection for the development of the pandemic is of yet not known, and should be proven with controlled trials of human responses to the bovine virus. One important outcome of this analysis is that if veterinary vaccines are to be considered for human use against COVID-19, as is being tested in Brazil, BCoV is a likely candidate.

## Conflict of interest

The authors declare no conflict of interest.

## Source of funding

This study was funded by the Coordenação de Aperfeiçoamento de Pessoal de Nível Superior (CAPES; Grant No 88881.505280/2020-01).

## Notes

### Competing Interest Statement

The authors have declared no competing interest.

